# Stator remodeling mechanism of *Bacillus subtilis* flagellar motor during biofilm development

**DOI:** 10.1101/2020.07.14.203471

**Authors:** Naoya Terahara, Keiichi Namba, Tohru Minamino

**Affiliations:** Graduate School of Frontier Bioscience, Osaka University, 1-3 Yamadaoka, Suita, Osaka 565-0871, Japan; Department of Physics, Faculty of Science and Engineering, Chuo University, 1-13-27 Kasuga, Bunkyo-ku, Tokyo 112-8551, Japan; RIKEN Spring-8 Center and Center for Biosystems Dynamics Research, 1-3 Yamadaoka, Suita, Osaka 565-0871, Japan; JEOL YOKOGUSHI Research Alliance Laboratories, Osaka University, 1-3 Yamadaoka, Suita, Osaka 565-0871, Japan

## Abstract

*Bacillus subtilis* possesses two distinct types of stator protein complexes for the flagellar motor: H^+^-type MotAB and Na^+^-type MotPS. The MotPS complex is used when both the external Na^+^ concentration and viscosity are high. Because deletion of the *motPS* genes does not affect swimming motility of cells, a physiological role of MotPS-dependent motility remains unclear. Here, we report that the MotPS stator complex is required for efficient biofilm maturation. Depletion of the MotPS complex did not cause a significant delay in the initiation of biofilm formation but reduced the number of viable cells in the biofilm. The MotAB stator units were efficiently replaced by the MotPS complexes with an increase in the viscosity of the environments. Therefore, we propose that MotPS-dependent motility of motile cells in the biofilm structure is required to efficiently keep the bacterial society in the biofilm healthy.

## INTRODUCTION

Planktonic bacterial cells can swim in liquids and move on solid surfaces by rotating long, helical filamentous structures called flagella. The flagellum is composed of the basal body acting as a rotary motor, the hook as a universal joint and the filament as a helical propeller^1,2^. Although the core structure of the flagella is highly conserved among bacterial species, novel and divergent structures surrounding the conserved structure have been revealed by in situ structural analyses of the flagella of different bacterial species, suggesting that each flagellum has evolved to function differently in various environments of the habitat of bacteria^3,4^.

The flagellar motor is composed of a rotor and multiple stator units, each of which acts as a transmembrane ion channel to couple the ion flow through the channel to torque generation caused by stator-rotor interactions^5,6^. The stator unit consists of three structural parts: a cytoplasmic domain responsible for the interaction with the rotor protein FliG, a transmembrane ion channel domain that form a pathway for the translocation of ions across the cytoplasmic membrane, and a peptidoglycan-binding (PGB) domain that anchors the stator unit to the peptidoglycan (PG) layer^7^. The flagellar motor of *Escherichia coli* and *Salmonella enterica* utilizes H^+^ as the coupling ion to generate the rotational force. In contrast, the flagellar motor of marine *Vibrio spp.* and extremely alkalophilic *Bacillus spp.* use Na^+^ as the coupling ion. Based on the coupling ion and primary sequence similarity, the stator units of the flagellar motor are divided into three groups: H^+^-type MotAB, Na^+^-type PomAB and Na^+^-type MotPS^7^. PomA and MotP are MotA homologues, and PomB and MotS are MotB homologues. Marine γ-proteobacteria, such as *Vibrio alginolyticus* and *Shewanella oneidensis* MR-1, have the Na^+^-type PomAB stator in the flagellar motor whereas extremely alkalophilic *Bacillus spp.* have the Na^+^-type MotPS stator^6^. Extremely alkalophilic *Bacillus clausii* encodes only the MotAB stator unit on the genome but this MotAB complex utilizes H^+^ at neutral pH and Na^+^ at extremely high pH for flagella-driven motility^8^. Furthermore, the MotPS stator unit of *Bacillus alcalophilus* Vedder 1934 conducts not only Na^+^ but also K^+^ to generate the rotational force^9^. Thus, each type of the stator unit appears to have been optimized for the best use of specific ions according to the environmental conditions.

Neutralophilic *Bacillus spp*. such as *Bacillus subtilis* have the Na^+^-coupled MotPS stator in addition to the H^+^-coupled MotAB stator^10^. The *B. subtilis* flagellar motor can accommodate about ten stator units around a rotor^11^. These two distinct stator complexes can be installed into the motor at the same time when external Na^+^ concentration is high^12,13^. The number of functional MotPS stator units in the motor increases from one to ten when the viscosity of the environment is increased by adding a polysaccharide, Ficoll 400, which is a neutral, highly branched, hydrophilic polysaccharide^11^. Recently, it has been shown that that the PGB domain of the MotPS stator acts as a viscosity sensor as well as a Na^+^ sensor^11,13^. Because deletion of the *motPS* genes does not affect the swimming motility of planktonic cells, the MotAB stator is dominant for wild-type motility under a wide range of experimental conditions^14^. Therefore, it remains unknown why neutralophilic *Bacillus spp.* have continued to maintain the *motPS* genes on the genome in addition to the *motAB* genes.

A chemical agent, 3’-5’ cyclic diguanylate monophosphate (c-di-GMP), is the second messenger that induces planktonic cells to form a sessile biofilm structure^15^. The *B. subtilis* flagellar motor is not only under the control of the sensory signal transduction pathway for chemotaxis but also under the control of the c-di-GMP signaling network^16^. Both EpsE and MotI suppress flagella-driven motility of *B. subtilis* planktonic cells, thereby leading to the motility-to-biofilm transition efficiently^17,18^. EpsE is encoded within the *eps* operon involved in the synthesis of extracellular polysaccharides, which are the compounds of the biofilm matrix^19^. EpsE directly binds to FliG to induce the dissociation of the MotAB stator units from the rotor^17,19^. MotI has a PilZ domain containing a conserved c-di-GMP binding motif. The c-di-GMP-bound form of MotI (MotI^c-di-GMP^) binds to MotA to inhibit flagellar motor rotation^18,20^. However, it remains unknown whether MotI^c-di-GMP^ binds to MotP like it does to MotA.

It has been reported that the MotPS stator complex also contributes to efficient biofilm formation^14^. However, it remains unknown how it does. To clarify this question, we investigated the impact of deletion of *motAB*, *motPS* or both stator genes on biofilm development and obtained novel insights into the stator remodeling mechanism of *B. subtilis* flagellar motor during the biofilm maturation process.

## RESULTS

### Effect of deletions of the *B. subtilis* stator genes on biofilm formation

Biofilm is a highly complex structure composed of extracellular polysaccharides, proteins, lipids and DNA. It has been observed that a subpopulation of planktonic *Bacillus* cells can swim in a biofilm structure by rotating flagella^21^. Because the number of active MotPS stator units in the motor increases from one to ten with an increase in the viscosity of the environments by adding Ficoll 400 in motor rotation assay media^11^, we hypothesized that the MotPS stator may play an important role in biofilm development. To clarify this hypothesis, we analyzed the capabilities of *B. subtilis* mutants with *motAB*, *motPS* and *motAB motPS* gene deletions to form biofilms. *B. subtilis* strain BR151MA, which is wild type for motility and chemotaxis, formed a floating membranous biofilm structure called pellicle at the air-water interface in a few days (Fig. 1a, 1st row), in agreement with a previous report^22^. The biofilm biomass of the BR151MA strain reached a maximum on day 5 and decreased thereafter (Fig. 1b, blue circle, and Supplementary Table 1). The MotAB strain (*motPS* deletion mutant) formed pellicles at the air-water interface in a similar manner (Fig. 1a, 2nd row). However, the biofilm biomass of the MotAB strain showed no reduction after day 5 (Fig. 1b, red triangle, and Supplementary Table 1). The MotPS strain (*motAB* deletion mutant) and stator-less strain (*motAB motPS* deletion mutant) also formed pellicles but showed a significant delay in pellicle formation (Fig. 1a, 3rd and 4th rows), indicating that the MotAB complex is required for efficient initiation of biofilm development whereas the MotPS complex is not required at all for biofilm formation. The biofilm biomass of the MotPS strain reached the wild-type level after 7 days of incubation (Fig. 1b, green square and Supplementary Table 1). Two-tailed *t*-test revealed no difference in the maximum biofilm biomass between the wild-type, MotAB and MotPS strains (Fig. 1c). The biofilm biomass of the stator-less strain reached the maximum level after 7 days of incubation (Fig. 1b, orange diamond and Supplementary Table 1) but it was slightly less than that of the MotPS cells (*P* < 0.05) (Fig. 1c). These observations suggest that flagella-driven motility of planktonic cells in the biofilm structure contributes to efficient maturation of the biofilm structure. The biofilm biomass of the MotPS strain decreased after day 8 in a similar manner to the wild-type BR151MA strain (Fig. 1b, green square). In contrast, the biofilm biomass of the stator-less strain showed almost no reduction like the MotAB strain (orange diamond), suggesting that MotPS-driven motility of planktonic cells in the biofilm structure is required for the decay of pellicles. Therefore, we propose that the MotPS stator contributes to an efficient transition from a sessile biofilm to planktonic motility.

**Fig. 1.**
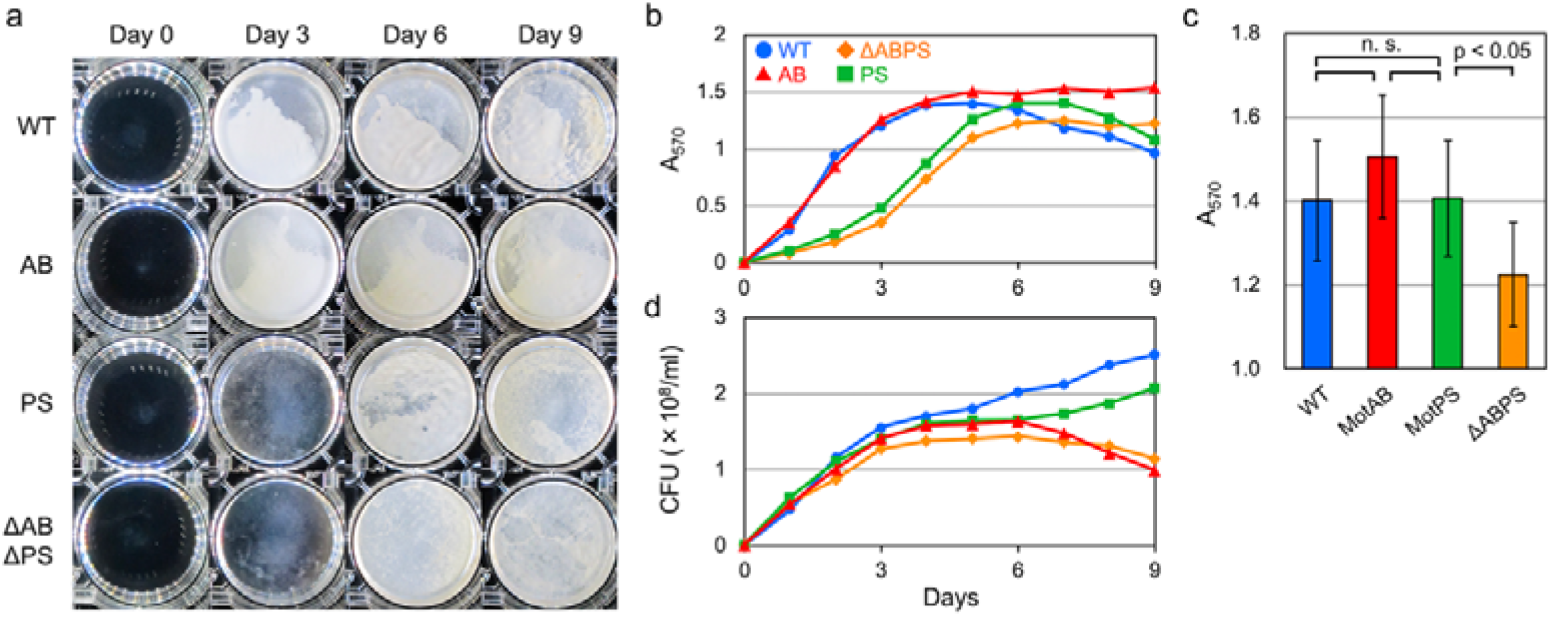
Effect of deletions of stator genes in biofilm formation. (a) Pellicle formation of the wild-type strain expressing both MotAB and MotPS complexes (indicated as WT), the MotAB strain (*motPS* deletion mutant, indicated as AB), the MotPS strain (*motAB* deletion mutant, indictated as PS) and the stator-less strain (*motAB motPS* deletion mutant, indicated as ΔABPS) at the air-water interface. (b) Measurements of the biofilm biomass by crystal violet assays. (c) Comparison of maximum biofilm biomass among the above strains. Paired *t*-tests showed a significant difference in the maximum biofilm biomass between the MotPS and ΔABPS strains (p < 0.05) but no significant difference (n. s.) among the WT, MotAB, MotPS strains. (d) Measurements of the number of viable cells in biofilms by plating the diluted culture onto nutrient agar plates.

### Effect of deletions of the *B. subtilis* stator genes on cell viability in a sessile biofilm structure

It has been reported that flagella-driven motility of planktonic cells in biofilms keeps the biofilm society healthy^21^. We found that the MotPS stator complex contributes to efficient dissolution of pellicles (Fig. 1b), leading to a plausible hypothesis that MotPS-driven motility of planktonic cells could contribute to cell viability in the pellicles. To test this possibility, we counted the number of viable cells of the wild-type, MotAB, MotPS and stator-less strains during the biofilm maturation process. The number of viable cells increased up to 3 days at almost the same rate in all four strains (Fig. 1d and Supplementary Table 1). The number of viable cells of the stator-less strain was almost constant after day 3 (Fig. 1d, orange diamond). The number of viable wild-type (blue circle) and MotPS (green square) cells gradually increased during biofilm development although the population of viable MotPS cells was slightly less than that of the wild-type. The number of viable MotAB cells slightly increased until day 6 in the same way to the MotPS cells but then decreased to the level of the stator-less strain (red triangle). These results suggest that the MotPS stator complex is required for cell viability in the biofilm society after prolonged incubation.

### Effects of EpsE and MotI on MotPS-dependent motility of planktonic cells

EpsE and MotI^c-di-GMP^ are responsible for an efficient motility-to-biofilm transition. These two proteins inhibit flagella-driven motility of planktonic cells, thereby initiating biofilm formation efficiently^16^. EpsE and MotI^c-di-GMP^ bind to FliG and MotA, respectively, thereby inhibiting torque generation caused by interactions between MotA and FliG^17–20^. We found that the MotPS strain showed a significant delay in biofilm formation (Fig. 1b), raising the possibility that the binding affinity of MotI for MotP could be lower than that for MotA. To clarify this possibility, we investigated how EpsE and MotI^c-di-GMP^ affect MotPS-dependent motility. The *motPS* genes are located downstream of the *ccpA* gene encoding a central regulator of carbon metabolism and form an operon along with the *ccpA* gene^23^. A stem-loop structure between the *ccpA* and *motP* genes serves as a transcriptional terminator of the *ccpA* gene, and hence the expression level of *motPS* is quite low although MotP and MotS are synthesized in the planktonic BR151MA cells^11–13^. Therefore, we used a *B. subtilis* strain expressing the *motP* and *motS* genes from the P_*motAB*_ promoter on the genome to increase the probability of MotPS stator assembly into the flagellar motor^11^. Over-expression of EpsE and MotI both inhibited free-swimming motility of the MotAB cells in soft agar (Fig. 2a), in agreement with previous reports^17,18^. The over-expression of EpsE inhibited MotPS-dependent motility in a similar manner (Fig. 2b). This is simply because EpsE binds to FliG. However, the over-expression of MotI did not affect MotPS-dependent motility (Fig. 2b). Therefore, we propose that a significant delay in biofilm formation of the MotPS cells may be a consequence of the inability of MotI to suppress the MotP function. However, because the swimming motility of the MotPS cells is about 5 times slower than that of the MotAB cells^12^, it is also possible that such a slow motile behavior of the MotPS strain is responsible for the delay in biofilm formation.

**Fig. 2.**
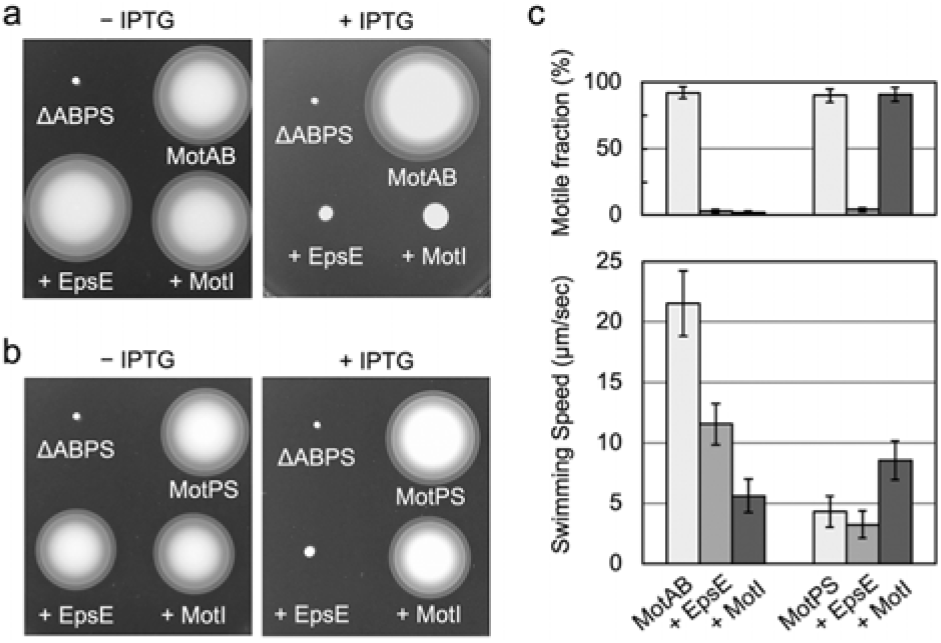
Multicopy effects of EpsE and MotI on flagella-driven motility of planktonic motile cells of *B. subtilis*. (a, b) Motility assays of the (a) MotAB and (b) MotPS strains transformed with a pHT01-based plasmid encoding EpsE or MotI in soft agar without (left panel) or with (right panel) IPTG. The stator-less strain (ΔABPS) was used as a negative control for motility assay. All plates were incubated at 37°C for 10 hours. (c) Multicopy effects of EpsE and MotI on free-swimming motility in liquid media. Motile fraction (upper panel) and the average swimming speed (lower panel) of motile cells were measured.

To test how many planktonic MotAB and MotPS cells over-expressing EpsE or MotI are motile, we next analyzed free-swimming motility in liquid media. Only a few % of MotAB cells over-expressing EpsE or MotI were motile (Fig. 2c, upper panel and Supplementary Table 2). The average speed of motile MotAB cells over-expressing EpsE was 11.6 ± 1.7 μm/sec, which was approximately half of the vector control (21.6 ± 2.7 μm/sec), and that of motile MotAB cells over-expressing MotI was 5.6 ± 1.4 μm/sec, which was about one fourth of the vector control (Fig. 2c, lower panel and Supplementary Table 2). Consistently, the size of motility ring on the soft agar of the MotAB cells over-expressing MotI was about twice larger than that of the MotAB cells over-expressing EpsE (Fig. 2a, right panel). On the other hand, only a few % of MotPS cells over-expressing EpsE were motile whereas more than 90% of cells over-expression of MotI were motile (Fig. 2c, upper panel and Supplementary Table 2). Over-expression of EpsE caused a slight decrease in the average swimming speed of motile MotPS cells from 4.3 ± 1.3 to 3.2 ± 1.1 μm/sec (Fig. 2c, lower panel and Supplementary Table 2). In contrast, the over-expression of MotI increased the average swimming speed by about 2-fold (8.6 ± 1.6 μm/sec) (Fig. 2c, lower panel and Supplementary Table 2), suggesting that MotI facilitates the MotPS motor activity significantly. Therefore, we propose that MotI^c-di-GMP^ may serve as a positive regulator to support the MotPS complex to efficiently assemble into the flagellar motor during the biofilm maturation process.

### Transcription levels of MotAB and MotPS stator genes during biofilm development

It has been shown that flagellar gene regulon is placed under control of c-di-GMP signaling networks so that flagellar gene transcription is suppressed during biofilm development^16^. The transcription level of the *motPS* genes is quite low in the planktonic motile cells due to a stem-loop structure between the *ccpA* and *motP* genes^12^. We found that the MotPS stator complex is required not only for cell viability in pellicles but also for efficient decay of the pellicles (Fig. 1d), raising the question of whether the transcription level of the *motPS* genes increases as biofilm matures. To investigate the transcription dynamics of the *motAB* and *motPS* genes in the biofilms, total RNA was extracted from pellicles of the BR151MA strain, and then the amount of each transcript was analyzed by RT-PCR (Fig. 3a and Supplementary Fig. 1). For first 4 days, the amount of the *motAB* mRNA decreased as the biofilm matured (Fig. 3b, filled circle). The *motAB* mRNA level decreased to 20% on day 4 as compared to the time of induction of biofilm formation and then increased again. In contrast, the *motPS* mRNA level increased as the biofilm matured, by a factor of 3 on day 5 compared to the start of culture (Fig. 3b, filled triangle). Interestingly, the *ccpA* mRNA level decreased as the biofilm matured (open triangle). Because the *motPS* genes are transcribed as an operon with the *ccpA* gene^23^, we assume that a conformational change in the stem-loop structure between the *ccpA* and *motP* genes may have occurred, which not only interferes with the termination of the *ccpA* transcripts but also stabilizes the *motPS* transcripts. Because we found that the biofilm biomass of the BR151MA strain began to decline on day 6 and that MotPS-dependent motility was required for efficient release of planktonic motile cells from pellicles (Fig. 1b), we suggest that such dynamic changes in the expression patterns of the *motAB* and *motPS* genes may contribute to efficient stator remodeling of the *B. subtilis* flagellar motor during the biofilm maturation process.

**Fig. 3.**
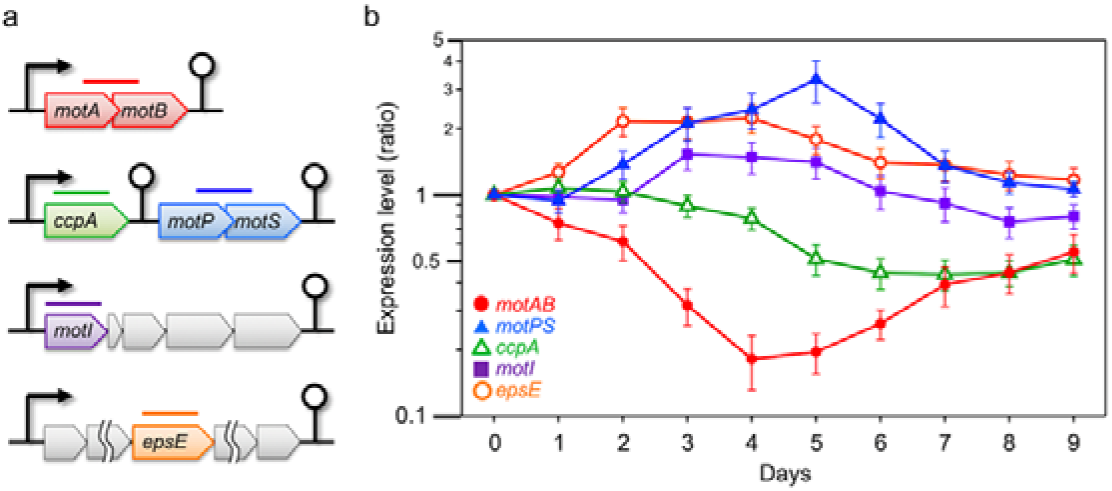
Measurements of the mRNA levels of stator genes during biofilm development (a) Amplified region (500 bp) in the target genes were shown by colored bar. (b) Relative transcription level of target genes during biofilm maturation. Each signal intensity of the EtBr-stained PCR product (Extended Data Fig. 1) was normalized to the initial value obtained in the start of culture. The transcription levels of each target gene were shown in logarithmic scale.

We found that MotAB-dependent motility is required for efficient initiation of biofilm formation (Fig. 1 b). The inhibition of MotAB-dependent motility by EpsE and MotI^c-di-GMP^ is required for efficient biofilm development^17–20^, raising the possibility that the expression levels of EpsE and MotI could be increased after induction of biofilm formation. The *epsE* mRNA levels increased after the start of culture and reached a maximum on day 2. Then, the *epsE* mRNA levels gradually decreased as the biofilm matured (Fig. 3b, open circle). The *motI* mRNA level was almost constant for the first 2 day but peaked on day 3. Then, the *motI* mRNA level gradually decreased as seen in the dynamics of the *epsE* mRNA level (filled square). Because we found that MotI facilitates MotPS-driven motility (Fig. 2), we propose that such dynamics of the *epsE* and *motI* gene expression are responsible not only for efficient motility-to-biofilm transition but also for efficient generation of MotPS-driven motile cells in the biofilm structure.

### Torque vs speed relationship of the wild-type *B. subtilis* flagellar motor under high-viscosity conditions

We found that MotPS-dependent motility is required for biofilm dispersion as well as cell viability in the biofilm structure after day 6 (Fig. 1). We also found that not only the *motPS* transcripts but also the *motAB* transcripts were detected in the biofilms after prolonged incubation (Fig. 3). Because the binding affinity of the MotPS complex for the motor is about 10-fold lower than that of the MotAB complex^11^, our present observations raise the question of how the MotPS stator units is more efficiently incorporated into the motor than the MotAB stator unit during the biofilm development process. Because the biofilm matrix is composed of extracellular polysaccharides, proteins and DNA, the viscosity of the biofilms should be very high. The number of active MotPS stator units in the motor increases from one to ten with an increase in the viscosity of the environment by adding Ficoll 400 in solution^11^. These raise the possibility that the environmental viscosity may also affect the binding affinity of the MotAB stator complex for the motor. To clarify this possibility, we analyzed the torque versus speed relationship of the MotAB motor under highly viscous conditions by bead assays (Fig. 4a and Supplementary Table 3).

**Fig. 4.**
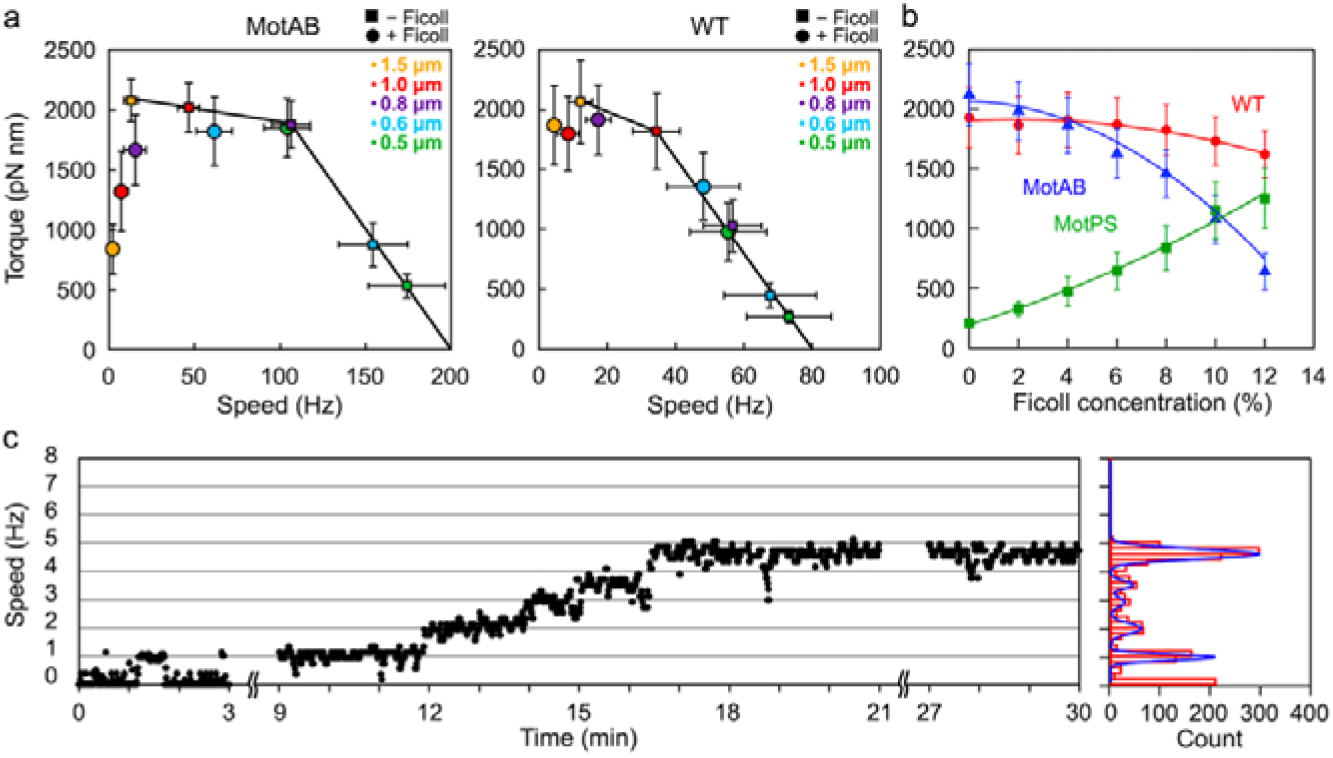
Rotation measurements of the *B. subtilis* flagellar motor over a wide range of external load. (a) Torque-speed curve of the MotAB (left panel) and wild-type (right panel) motors in the presence and absence of 10%(w/v) Ficoll-400. Rotation measurements were carried out at 23°C by tracking the position of 1.5-μm (orange), 1.0-μm (red), 0.8-μm (purple), 0.6-μm (cyan) and 0.5-μm (light green) beads attached to the partially sheared sticky filament of the flagellar motor. More than twenty individual beads of each size were measured. (b) Torque of the wild-type (red), MotAB (blue) and MotPS (green) motors measured using a 1.0-μm bead in media containing 0%, 2%, 4%, 6%, 8% and 10% Ficoll 400 (w/v). (c) Speed increment in a single MotAB motor by resurrection experiment. The trace of speed was measured using a 1.0 μm bead in the presence of 10% Ficoll (left panel), and then size of speed increment was determined by histogram analysis and multiple Gaussian fitting (right panel).

The torque-speed curve of the flagellar motor is composed of a high-load, low-speed regime and a low-load, high-speed regime. Torque is almost constant in the high-load, low-speed regime but decreases steeply in the low-load, high-speed regime^24^. The *B. subtilis* MotAB motor showed a typical torque-speed curve in the absence of Ficoll 400 (Fig. 4a, left panel, square), in agreement with a previous report^11^. However, the MotAB motor showed an unusual torque-speed curve in the presence of 10% Ficoll 400 (circle). The maximum torque produced by the MotAB motor decreased from about 2,081 ± 174 pN nm to about 837 ± 210 pN nm with an increase in the viscosity of the environments by adding 10% Ficoll 400 (Supplementary Table 3), indicating that a large increment in the viscosity considerably inhibits the MotAB motor activity. In contrast, the addition of Ficoll 400 to the motility buffer did not reduce the maximum torque produced by the wild-type motor (Fig. 4a, right panel and Supplementary Table 4), suggesting that the wild-type motor rapidly exchanges between the MotAB and MotPS stator units around the rotor in response to changes in the viscosity of the environment. To confirm this, we measured the rotation speed of the wild-type, MotAB and MotPS motors labelled with a single 1.0 μm bead in motility buffers containing Ficoll 400 over a range of concentrations from 0% to 10% (w/v) in the presence of 200 mM NaCl (Supplementary Table 5). As the concentration of Ficoll 400 was increased step-by-step by 2% at each step, torque produced by the MotPS motor increased (Fig. 4b, green square), in agreement with a previous report^11^. However, torque produced by the MotAB motor decreased with an increase in the Ficoll concentration (blue triangle), and the wild-type motor torque was almost constant (red circle). The maximum torque is dependent on the number of active stator units in the *B. subtilis* flagellar motor^11^ as seen in the *E. coli* and *S. enterica* flagellar motors^25–27^. As a single MotPS stator unit can produce nearly the same torque as a single MotAB stator unit^11^, these results lead to a plausible hypothesis that the binding affinity of the MotAB stator unit for the motor decreases as the viscosity of the environment increases, unlike that of the MotPS complex, which increases as the viscosity increases.

The *B. subtilis* flagellar motor can accommodate ten active stator units around a rotor^11^. To quantitatively determine the number of active stators in the *B. subtilis* MotAB motor under highly viscous conditions, we carried out resurrection experiments in the presence of Ficoll 400 to measure the stepwise increment in the rotation rate upon induction of the MotAB stator from the IPTG-inducible *P*_*grac*_ promoter. The speed increment in a single MotAB motor labelled with a 1.0-μm bead is shown in Fig. 4c. The unit increment was 0.9 Hz as judged by multiple Gaussian fitting of speed histograms (right panel), and the torque produced by a single MotAB stator unit was calculated to be 203.8 ± 34.8 pN nm in the presence of 10% Ficoll 400, which was nearly the same torque value obtained in the absence of Ficoll^11^. This indicates that Ficoll 400 does not directly inhibit the MotAB stator function. Because each incremental unit in the motor speed reflects the incorporation of a single stator unit around the rotor, the number of active MotAB stator units in the motor labeled with the 1.0-μm bead decreased from 10 to 5 by adding 10% Ficoll 400. These results suggest that the MotAB and MotPS stator units autonomously modulates their binding affinities for the motor in response to changes in the viscosity of the environments to maintain the optimal motor performance under various environmental conditions.

## DISCUSSION

The flagellar motor can detect changes in the environments, not only coordinating motor activity in response to the environmental changes but also inducing a motility-to-biofilm transition^3,7^. *B. subtilis* encodes two distinct types of stators on the genome: H^+^-type MotAB and Na^+^-type MotPS^10^. It has been shown that the MotAB stator complex is dominant for free-swimming motility of planktonic cells under various experimental conditions^14^. Deletion of the *motPS* genes has no impact on the swimming motility although the MotPS stator complex is expressed in planktonic cells^12,28^. It has been reported that increases in both external Na^+^ concentrations and the environmental viscosity significantly improve the assembly efficiency of the MotPS stator complex into the motor^11,13^. Since the biofilm matrix is composed of extracellular polysaccharides, proteins, DNA and lipids, all of which increase the viscosity, it leads to a plausible hypothesis that MotPS-dependent motility could contribute to biofilm development^11^.

To clarify this hypothesis, we analyzed the capabilities of the wild-type, MotAB and MotPS strains to form biofilms (Fig. 1). The MotPS strain formed a floating membranous biofilm structure as seen in the wild-type and MotAB strains (Fig. 1a). However, the MotPS strain exhibited a significant delay in biofilm assembly compared to the wild-type and MotAB strains (Fig. 1b), indicating that MotAB-dependent motility contributes to efficient initiation of biofilm formation. The biofilm biomass of the wild-type and MotPS strains decreased after a prolonged incubation whereas that of the MotAB strain was almost constant (Fig. 1b), indicating that MotPS-dependent motility is required for efficient decay of the biofilm to induce a biofilm-to-motility transition. Furthermore, the viable cell populations increased in the wild-type and MotPS biofilms but decreased in the MotAB biofilms (Fig. 1d), suggesting that MotPS-dependent motility of motile cells in the biofilms is important in keeping the biofilm society healthy. Therefore, the reason why neutralophilic *Bacillus* species have continued to maintain the *motPS* genes on the genome throughout the evolutional process may be for such an efficient cell survival strategy within biofilms.

EpsE and MotI^c-di-GMP^ bind to FliG and MotA, respectively, inhibiting flagellar motor rotation to induce biofilm formation efficiently^17,18^. Here, we found that over-expression of MotI inhibited MotAB-dependent motility but not MotPS-dependent motility (Fig. 2). Interestingly, the over-expression of MotI increased the swimming speed of the motile MotPS cells by about 2-fold (Fig. 2c), indicating that MotI act as a positive regulator of the MotPS-dependent motility. It has been shown that the swimming speed of motile cells depends on the number of active stator units in the motor^29,30^. Because the MotPS motor has only a single stator unit around a rotor in planktonic motile MotPS cells even when external Na^+^ concentration is high^11^, we propose that the interaction between MotP and MotI may increase the binding affinity of the MotPS stator complex for the motor, thereby increasing the stator number in the MotPS motor even in the planktonic cells. We are currently testing this hypothesis.

The levels of *motPS* transcripts are relatively low in planktonic *B. subtilis* cells because of a stem-loop structure between the *ccpA* and *motP* genes^12^. Here, we found that the levels of the *motPS* transcripts increased by a factor of 3 as the biofilm matured (Fig. 3) whereas the mRNA levels of the *motAB* genes decreased to 20%. Therefore, we propose that such dynamic changes in the transcription levels of the *motAB* and *motPS* genes allow the MotPS stator complex to efficiently work in the flagellar motor of motile cells in the biofilm structure.

How does the *B. subtilis* flagellar motor efficiently exchange the MotAB and MotPS stator complexes in response to changes in lifestyle? The flagellar motor autonomously controls not only its ion channel activity^31^ but also the number of active stator units in response to changes in the viscosity of the environment^27,32^. It has been reported that certain mutations in the cytoplasmic domain of the MotAB complex and an in-frame deletion in the linker region of the MotAB complex connecting the transmembrane H^+^ channel domain and the PGB domain affect the mechano-sensitivity of the flagellar motor, thereby showing an unusual torque-speed curve^26,33^. These observations suggest that the cytoplasmic domain of the stator complex acts as a load sensor to control load-dependent assembly–disassembly dynamics of the stator complex and that the flexible linker region regulates the binding affinity of the PGB domain for the PG layer in an external-load dependent manner^26,33^. The PGB domain of the MotPS stator complex acts as a viscosity sensor as well as a Na^+^ sensor, and hence the number of active MotPS stator units increases from one to ten at an elevated external Na^+^ concentration and viscosity^11,13^. Here, we showed by single motor bead assay that the number of active stator units in the MotAB motor decreased from 10 to 5 with an increase in the viscosity by adding Ficoll 400 in the motility buffer, thereby reducing the maximum torque by 50%. However, the torque produced by the wild-type motor was almost constant, suggesting that rapid exchanges between the MotAB and MotPS stator units occur in the wild-type motor in response to changes in the viscosity of the environment. Therefore, we propose that the MotAB and MotPS stator complexes autonomously modulate their binding affinity for the PG layer in response to the viscosity changes.

Based on the available information, we propose a plausible stator remodeling mechanism of the *B. subtilis* flagellar motor (Fig. 5). Planktonic motile cells predominantly use the H^+^-type MotAB complex as an active stator unit in the motor. When the motile cells attach to the solid surface or reach the water-air interface, both EpsE and MotI^c-di-GMP^ inhibit MotAB-dependent motility, thereby inducing biofilm formation efficiently. As the biofilm matures, the levels of *motPS* transcripts increase whereas those of the *motAB* transcripts decrease. Furthermore, the viscosity is also increased in the biofilm structure due to the accumulation of huge amounts of extracellular polysaccharides. Because intracellular c-di-GMP concentration of the cells living in the biofilm structure is relatively high, MotI^c-di-GMP^ facilitates the assembly of the MotPS complex into the flagellar motor but inhibits the assembly of the MotAB complex. As a result, the flagellar motor can efficiently accommodate multiple MotPS stator units around the rotor to generate MotPS-dependent motility in biofilms, thereby not only keeping the biofilm society healthy but also inducing the biofilm-to-motility transition.

**Fig. 5.**
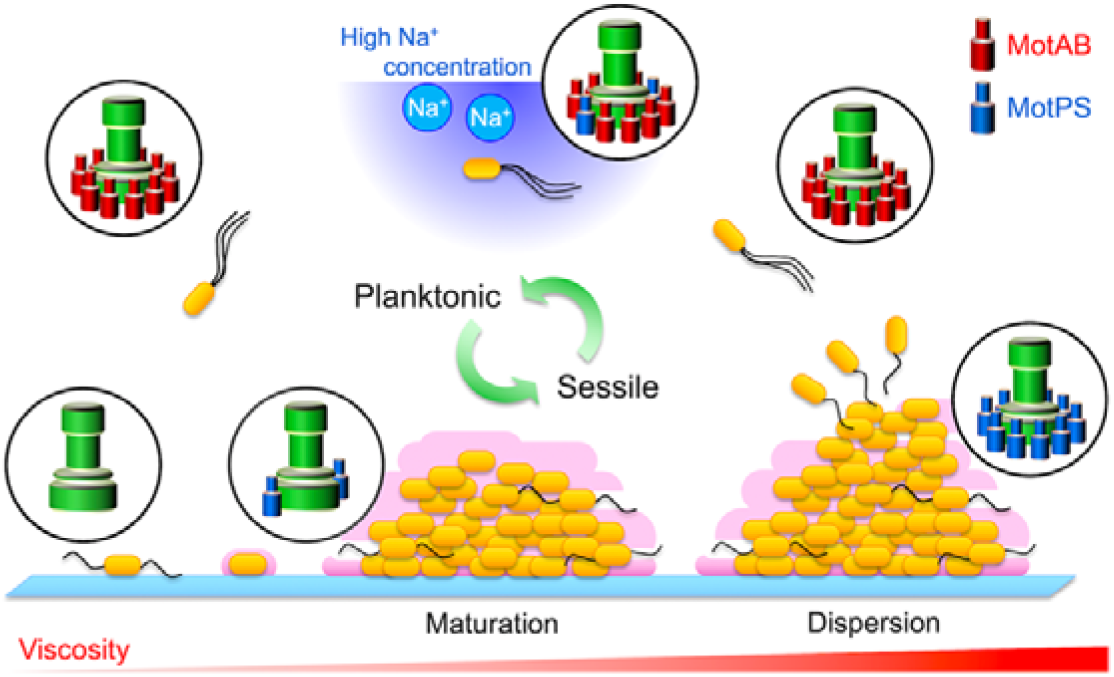
Stator remodeling mechanism of the *B. subtilis* flagellar motor during lifecycle. Planktonic *B. subtilis* cells predominantly use the H^+^-type MotAB complex (red) as a functionally active stator unit in the flagellar motor. When planktonic cells attach to solid surfaces or reach the water-air interface, c-di-GMP signaling networks induce a motility-to-biofilm transition. EpsE and MotI^c-di-GMP^ inhibit MotAB-dependent motility, thereby promoting the biofilm formation efficiently. The transcription level of the *motPS* genes increases as the biofilm matures whereas that of the *motAB* genes decreases. During biofilm maturation, the viscosity is increased in the biofilm structure due to the accumulations of extracellular polysaccharides, protein, DNA and so on, and hence the binding affinity of the MotPS stator unit (blue) for the motor increases considerably whereas that of the MotAB stator complex decreases significantly. Furthermore, MotI^c-di-GMP^ facilitates the assembly of the MotPS complex into the motor but suppresses that of the MotAB complex. As a result, the flagellar motor can predominantly accommodate multiple MotPS stator units around the rotor during biofilm development. MotPS-driven motility of a very small population of planktonic cells in the biofilms not only keeps the biofilm society healthy but also efficiently induces the biofilm-to-motility transition. After dissociation of planktonic cells from the biofilm, the binding affinity of the MotPS complex for the motor decreases whereas that of the MotAB complex increases, allowing planktonic motile cells to predominantly use the MotAB complex as a stator unit of the motor again for rapid and efficient movements towards more suitable conditions.

## METHODS

### Bacterial strains, plasmids and media

Bacterial strains and plasmids used in this study are listed in Supplementary Table 6. L-broth (LB) contained 1% Bacto tryptone, 0.5% yeast extract, 0.5% NaCl. LB soft agar plates contained 1% Bacto tryptone, 0.5% yeast extract, 0.5% NaCl, 0.25% Bacto agar with or without 0.6 mM IPTG. MSgg medium (pH 7.0) contained 5 mM K_2_HPO_4_, 100 mM MOPS, 2 mM MgCl_2_, 700 μM CaCl_2_, 50 μM MnCl_2_, 50 μM FeCl_3_, 1 μM ZnCl_2_, 2 μM thiamine, 0.5% glycerol, 0.5% glutamate, 50 μg/ml tryptophan and 50 μg/ml lysine^22^. Ampicillin, erythromycin, neomycin, chloramphenicol and spectinomycin were added to the medium at a final concentration of 50 μg/ml, 0.3 μg/ml, 7.5 μg/ml, 5 μg/ml and 150 μg/ml, respectively, if necessary.

### DNA manipulations

All primers used for plasmid construction are listed in Supplementary Table 7. DNA manipulations were carried out as described previously^11^. DNA sequencing reactions were carried out using BigDye v3.1 (Applied Biosystems) and then the reaction mixtures were analyzed by a 3130 Genetic Analyzer (Applied Biosystems).

### Motility assays

Fresh colonies were inoculated onto 0.25% LB soft agar plates, and the plates were incubated at 37°C for 10 hours. To measure the swimming speed of *B. subtilis* cells, the cells were grown in 1.5 ml LB with shaking at 37°C until the cell density has reached an optical density of 1.0 at 600 nm and then placed under a phase contrast microscope. The swimming speed analysis and swimming cell counting were carried out using Move-tr/2D software (Library Co., Ltd.). More than 20 motile cells were analyzed in each measurement. Data were the average of three independent measurements.

### Biofilm formation assays

*B. subtilis* cells were grown overnight in 5 ml LB with shaking at 37°C. 40 μl of the overnight cultures were inoculated into 4 ml MSgg medium in 6-well plates, and pellicle biofilms were grown at 25°C. To quantify the biofilm biomass, the biofilms were stained by a dye, crystal violet, as described previously^34^. The biofilms were harvested by centrifugation, resuspended in 1 ml of 0.1% crystal violet solution and incubated for 20 min at room temperature. After washing twice with 1 ml of pure water to remove free dyes, the dyes attached to the biofilms were extracted in 1 ml of 95% EtOH for 10 min. After centrifugation to remove the biofilms, the absorbance of each supernatant was measured at 590 nm. To count viable cells in the biofilm, the biofilms were harvested by centrifugation, resuspended and diluted into a distilled water. The viable cells in the suspensions were determined by counting the number of colonies grown on LB agar plates. All data were the averages of three independent experiments.

### Statistical analysis

Statistical analyses were done using StatPlus::mac software (AnalystSoft). Comparisons were performed using a two-tailed Student’s *t*-test. A *P* value of < 0.05 was considered to be statistically significant difference.

### Reverse transcription PCR analysis

The total RNA was isolated from biofilm matrix using RNeasy PowerBiofilm Kit (QIAGEN). A 1.5 μg of total RNA was applied to reverse-transcription reaction for cDNA by using High Capacity RNA-to-cDNA Kit (Applied Biosystems). All specific primers for the target genes are listed in Table S2. The parts of the *motAB*, *motPS*, *ccpA*, *motI* and *epsE* genes were amplified by PCR using cDNA as the template. 5S rRNA was amplified using rrn-RT-F and rrn-RT-R primers as an internal control. PCR products were resolved by agarose gel electrophoresis and visualized by ethidium bromide. Each band intensity was analyzed by ImageJ version 1.48 (National Institutes of Health). The expression level of each gene was normalized to the values obtained in day 0. All data were the averages of three independent experiments.

### Bead assays

Bead assays for the *B. subtilis* flagellar motor were carried out in a motility buffer (10 mM potassium phosphate, pH 7.0, 0.1 mM EDTA, 10 mM lactate, 200 mM NaCl) as described previously^11^. Polystyrene beads with a diameter of 1.5 μm, 1.0 μm, 0.8 μm, 0.6 μm or 0.5 μm (Invitrogen) were used. Phase-contrast images of each bead were captured by a high-speed camera (Digimo-VCC-SXGA-B, Digimo) at the frame rate of 1,000 frames per second. The rotation speed analysis and torque calculation were carried out as described previously^35^. In resurrection experiments, *B. subtilis* cells were incubated in fresh LB containing 0.1 mM IPTG for 20 min 37°C with shaking. Then, bead assays using 1.0 μm beads were carried out in the motility buffer containing 1.0 mM IPTG and 10% Ficoll 400 as described previously^11^.

## DATA AVAILBILITY

All data generated during this study are included in this published article and its Supplementary Information files.

## ACKNOWLEDGEMENTS

This research has been supported in part by the Japan Society for the Promotion of Science (JSPS KAKENHI Grant Numbers, JP18K06085 to N.T., JP25000013 to K.N. and JP19H03182 to T.M.), JEOL YOKOGUSHI Research Alliance Laboratories of Osaka University to K.N and a grant from the Institute of Fermentation, Osaka, Japan to T.M.

## AUTHOR CONTRIBUTIONS

N.T., K.N. and T.M. conceived and designed the project. N.T. preformed the experiments; N.T. analyzed the data. N.T. wrote the paper. K.N. and T.M. supervised the study and edited the paper. All author reviewed and approved the paper.

