## Supplementary information for "Stator remodeling mechanism of *Bacillus subtilis* flagellar motor during biofilm development"

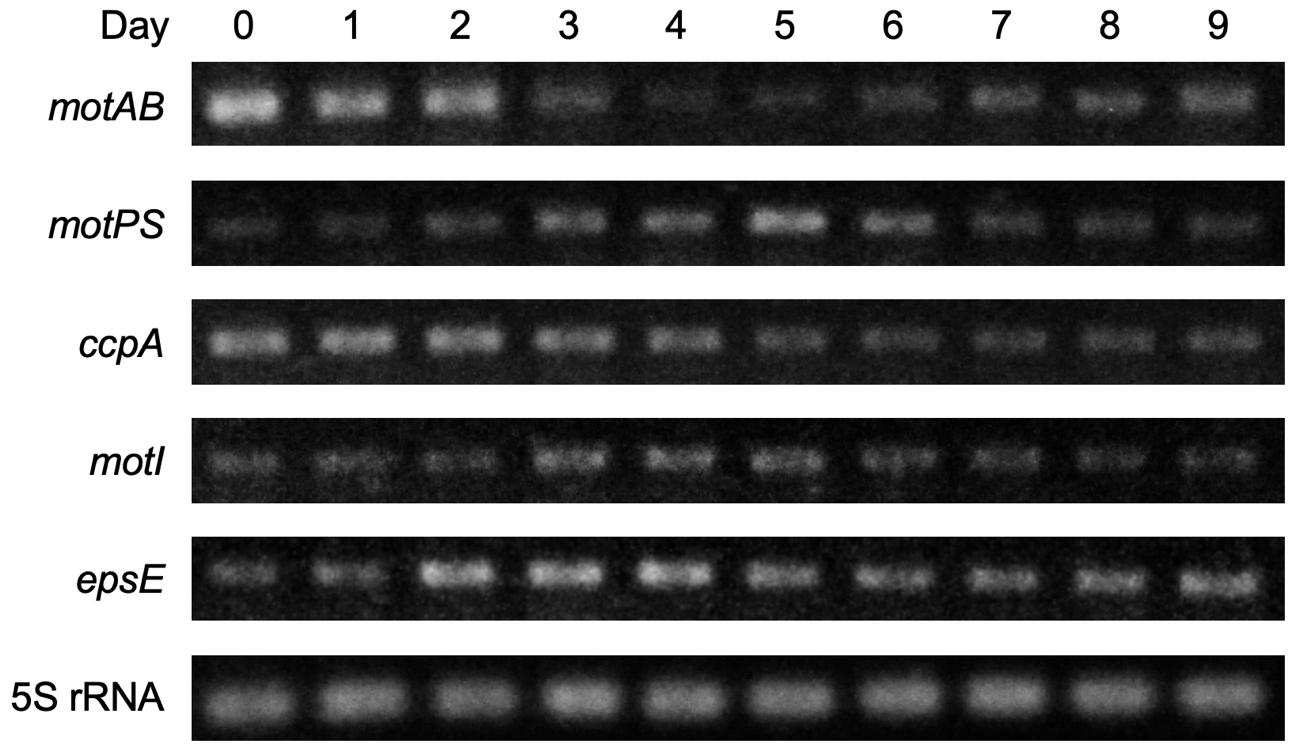

**Supplementary Fig. 1.** RT-PCR measurements of the transcripts of the stator genes during biofilm development. Target genes were amplified from cDNA. 5S rRNA was amplified as an internal control. The PCR products were subjected to agarose gel electrophoresis, followed by staining with ethidium bromide.

**Supplementary Table 1. Biomass and viable cell number during biofilm development**

|  | Days | 0 | 1 | 2 | 3 | 4 | 5 | 6 | 7 | 8 | 9 |
| --- | --- | --- | --- | --- | --- | --- | --- | --- | --- | --- | --- |
| WT | A_570_ | 0.005 ± 0.001 | 0.292 ± 0.039 | 0.945 ± 0.111 | 1.212 ± 0.132 | 1.384 ± 0.141 | 1.402 ± 0.143 | 1.341 ± 0.138 | 1.186 ± 0.124 | 1.113 ± 0.121 | 0.965 ± 0.118 |
|  | CFU  (×10^8^/ml) | 0.012 ± 0.0006 | 0.48 ± 0.004 | 1.16 ± 0.06 | 1.56 ± 0.09 | 1.71 ± 0.13 | 1.81 ± 0.16 | 2.03 ± 0.21 | 2.12 ± 0.23 | 2.38 ± 0.24 | 2.51 ± 0.28 |
| ΔAB  ΔPS | A_570_ | 0.007 ± 0.001 | 0.089 ± 0.012 | 0.189 ± 0.022 | 0.356 ± 0.043 | 0.747 ± 0.106 | 1.102 ± 0.114 | 1.227 ± 0.127 | 1.225 ± 0.124 | 1.203 ± 0.122 | 1.224 ± 0.122 |
|  | CFU  (×10^8^/ml) | 0.014 ± 0.0008 | 0.57 ± 0.007 | 0.88 ± 0.022 | 1.28 ± 0.07 | 1.38 ± 0.10 | 1.41 ± 0.11 | 1.44 ± 0.12 | 1.36 ± 0.11 | 1.31 ± 0.10 | 1.15 ± 0.08 |
| MotAB | A_570_ | 0.006 ± 0.001 | 0.362 ± 0.033 | 0.852 ± 0.104 | 1.263 ± 0.123 | 1.423 ± 0.141 | 1.506 ± 0.146 | 1.477 ± 0.141 | 1.533 ± 0.139 | 1.501 ± 0.137 | 1.543 ± 0.148 |
|  | CFU  (×10^8^/ml) | 0.018 ± 0.0009 | 0.55 ± 0.007 | 1.02 ± 0.04 | 1.42 ± 0.08 | 1.58 ± 0.11 | 1.60 ± 0.12 | 1.64 ± 0.13 | 1.48 ± 0.09 | 1.22 ± 0.07 | 1.00 ± 0.04 |
| MotPS | A_570_ | 0.006 ± 0.001 | 0.114 ± 0.016 | 0.254 ± 0.034 | 0.484 ± 0.052 | 0.884 ± 0.111 | 1.267 ± 0.121 | 1.404 ± 0.139 | 1.407 ± 0.138 | 1.279 ± 0.120 | 1.082 ± 0.118 |
|  | CFU  (×10^8^/ml) | 0.017 ± 0.0007 | 0.64 ± 0.008 | 1.11 ± 0.05 | 1.41 ± 0.08 | 1.61 ± 0.12 | 1.65 ± 0.12 | 1.66 ± 0.12 | 1.74 ± 0.18 | 1.88 ± 0.22 | 2.08 ± 0.24 |

**Supplementary Table 2. Swimming speed and motile fraction**

|  |  |  | + EpsE | + MotI |
| --- | --- | --- | --- | --- |
| MotAB | Speed (μm/sec) | 21.6 ± 2.7 | 11.6 ± 1.7 | 5.6 ± 1.4 |
|  | Fraction (%) | 92.3 ± 4.4 | 3.2 ± 1.5 | 2.1 ± 1.3 |
| MotPS | Speed (μm/sec) | 4.3 ± 1.3 | 3.2 ± 1.1 | 8.6 ± 1.6 |
|  | Fraction (%) | 90.4 ± 5.1 | 4.2 ± 1.6 | 91.1 ± 5.2 |

**Supplementary Table 3. Rotational speed and torque of the MotAB motor.**

| Ficoll | Bead size (μm) | 1.5 | 1.0 | 0.8 | 0.6 | 0.5 |
| --- | --- | --- | --- | --- | --- | --- |
| 0% | Speed (Hz) | 13 ± 2 | 47 ± 6 | 107 ± 15 | 155 ± 20 | 175 ± 33 |
|  | Torque (pN nm) | 2,081 ± 174 | 2,021 ± 205 | 1,879 ± 194 | 881 ± 182 | 535 ± 99 |
| 10% | Speed (Hz) | 2 ± 1 | 7 ± 4 | 15 ± 6 | 62 ± 10 | 104 ± 14 |
|  | Torque (pN nm) | 837 ± 210 | 1,320 ± 332 | 1,664 ± 294 | 1,820 ± 287 | 1,850 ± 243 |

**Supplementary Table 4. Rotational speed and torque of the wild-type motor.**

| Ficoll | Bead size (μm) | 1.5 | 1.0 | 0.8 | 0.6 | 0.5 |
| --- | --- | --- | --- | --- | --- | --- |
| 0% | Speed (Hz) | 12 ± 4 | 34 ± 7 | 57 ± 9 | 68 ± 14 | 73 ± 12 |
|  | Torque (pN nm) | 2,062 ± 348 | 1,817 ± 316 | 1,029 ± 216 | 450 ± 105 | 270 ± 52 |
| 10% | Speed (Hz) | 5 ± 2 | 9 ± 3 | 17 ± 4 | 48 ± 11 | 55 ± 12 |
|  | Torque (pN nm) | 1,865 ± 330 | 1,798 ± 310 | 1,912 ± 294 | 1,356 ± 280 | 977 ± 241 |

**Supplementary Table 5. Measurements of torque produced by the wild-type (WT), MotAB and MotPS motors using a 1.0-μm bead in media containing 0%, 2%, 4%, 6%, 8%, 10% and 12% Ficoll 400 (w/v).**

| Ficoll (%) | 0 | 2 | 4 | 6 | 8 | 10 | 12 |
| --- | --- | --- | --- | --- | --- | --- | --- |
| WT | 1,930 ± 253 | 1,866 ± 243 | 1,910 ± 231 | 1,871 ± 223 | 1,826 ± 212 | 1,732 ± 205 | 1,622 ± 198 |
| MotAB | 2,122 ± 262 | 1,984 ± 246 | 1,864 ± 233 | 1,628 ± 208 | 1,559 ± 199 | 1,278 ± 203 | 641 ± 153 |
| MotPS | 209 ± 35 | 331 ± 62 | 473 ± 124 | 645 ± 156 | 838 ± 185 | 1,150 ± 238 | 1,252 ± 251 |

**Supplementary Table 6. Bacterial strains and plasmids used in this study**

| Strain | Relevant characteristics | Source or reference |
| --- | --- | --- |
| *Escherichia coli* |  |  |
| DH5αMCR | F^－^ *mcrA*Δ*1* (*mrr-hsd RMS-mcrBC*) Ф80*dlacZ* Δ(*lacZYAargF*) *U169 deoR recA1 endA1 supE44 λthi-1 gyr-496 relA1* | Stratagene |
| *Bacillus subtilis* |  |  |
| BR151MA | *lys3 trpC2* (wild type for motility and chemotaxis) | 1 |
| ΔAB | *lys3 trpC2* Δ*motAB*::*ery* | 2 |
| ΔPS | *lys3 trpC2* Δ*motPS*::*neo* | 2 |
| ΔABΔPS | *lys3 trpC2* Δ*motAB*::*ery* Δ*motPS*::*neo* | 2 |
| AB | ΔABΔPS *amyE*::P*_motAB_*-*motAB* | 2 |
| PS | ΔABΔPS *amyE*::P*_motAB_*-*motPS* | 2 |
| ΔABΔPSΔHag | ΔABΔPS Δ*hag*::*spec* | 3 |
| AB-sticky | ΔABΔPSΔHag *amyE*::P*_motAB_*-*motAB*, P*_hag_*-*hagsticky* | 3 |
| PS-sticky | ΔABΔPSΔHag *amyE*::P*_motAB_*-*motPS*, P*_hag_*-*hagsticky* | 3 |
| P_grac_-AB-sticky | ΔABΔPSΔHag *amyE*::P*_grac_*-*motAB*, P*_hag_*-*hagsticky* | 3 |
| P_grac_-PS-sticky | ΔABΔPSΔHag *amyE*::P*_grac_*-*motPS*, P*_hag_*-*hagsticky* | 3 |
| Plasmid | Relevant characteristics | Source or reference |
| pHT01 | *B. subtilis* expression vector by P*_grac_* promoter | Mo Bi Tec |
| pHT-*epsE* | pHT01 + *epsE* | This study |
| pHT-*motI* | pHT01 + *motI* | This study |

**Supplementary Table 7. Oligonucleotides used in this study**

| Primer | Sequence (5’ → 3’) |
| --- | --- |
| epsE-BamHI-F | cgggatccATGAACTCAGGACCAAAAGTTTC |
| epsE-XmaI-R | tccccccgggCTTGACAAGCCCTTCCTTTTGGT |
| motI-BamHI-F | cgggatccATGATAGAGATTGGAGAAAATGT |
| motI-XmaI-R | tccccccgggTTATTCCATTCGGGCCTTTCTTC |
| ccpA-RT-F | GCGCGCGGAATTGAAGATATCGCG |
| ccpA-RT-R | CGAGTGCCATTTCATCAGTTGCAG |
| motPS-RT-F | TCGGACCAAACATGGCCATTGCGC |
| motPS-RT-R | CTTGCTTCTTCGTATCAGATGGGG |
| motAB-RT-F | TGCCTTTGTTGCCACACTTCTCGG |
| motAB-RT-R | TCACTTCATCGATGCCGTCTGACT |
| motI-RT-F | TTGAAAAAGGCAAAAAGCAAGGCG |
| motI-RT-R | GCAAAGCCTGCTGGTCACCTGCTG |
| epsE-RT-F | ACAGACGGCACGCTCCGTATCGCG |
| epsE-RT-R | AGGAAGCTTCAAGCGTCTGCACGC |
| rrn-RT-F | TGGTGGCGATAGCGAAGAGGTCAC |
| rrn-RT-R | TGGCGGCGTCCTACTCTCACAGGG |

Extra nucleotides that were added to introduce restriction sites are shown by a small letter.
